# Neutrophil FcγRI expression as a determinant of oxidative responses in human blood

**DOI:** 10.1101/2025.10.01.679808

**Authors:** Sandrine Huot, Paul R. Fortin, Cynthia Laflamme, Marc Pouliot

## Abstract

Neutrophils express Fc receptors on their surface to trap immune complexes. While the roles of FcγRIIa and FcγRIIIb have extensively been studied in that context, that of FcγRI remains elusive. Recently, aggregated IgGs have been shown to induce rapid FcγRI up-regulation and reactive oxygen species (ROS) generation, but the biological relevance of this process is still unclear. In this study, incubation of blood samples from healthy volunteers with heat-aggregated IgGs, used as a model of immune complexes, rapidly up-regulated the surface expression of FcγRI, predominantly on neutrophils, as measured by flow cytometry. Stimulation of isolated neutrophils with aggregated IgGs resulted in the production of ROS in an FcγRI-dependent fashion, as monitored with a luminol-based chemiluminescence assay. Cytochalasin B potentiated FcγRI expression and ROS production. In resting blood, positive correlations between neutrophil FcγRI and ROS production were observed, both in healthy volunteers and patients with lupus. This study unveils a potentially central regulatory role for neutrophil FcγRI in ROS production, both in healthy individuals and patients with lupus, and identifies neutrophil FcγRI as a promising target to modulate oxidative response.

## INTRODUCTION

Neutrophils, the most abundant leukocyte population in circulation, play central roles in host defence by eliminating immunoglobulin-coated microbes and tissue debris [1]. Through their Fc receptors, neutrophils recognize immune complexes and, in response, can engage a range of effector functions including phagocytosis, production of reactive oxygen species (ROS), release of lipid mediators, and neutrophil extracellular traps (NETs) [2, 3]. Human neutrophils express several Fcγ receptors (FcγRs) including the low-affinity FcγRIIa (CD32a) and FcγRIIIb (CD16b), with specialized, context-dependent functions [4–6]. FcγRIIa is a critical mediator of antibody-driven inflammation in autoimmunity [5], and is required for neutrophil recruitment within glomerular capillaries following IgG deposition [7, 8]. Its intracellular domain contains an immunoreceptor tyrosine-based activation motif (ITAM) which, upon activation, can trigger downstream signalling pathways to promote phagocytosis, ROS production, and cytokine secretion [9]. FcγRIIIb, unique to neutrophils, is anchored by a glycosylphosphatidylinositol (GPI) linker and lacks a cytoplasmic domain [10], relying on partners for signal transduction. FcγRIIIa (CD16a), a canonical ITAM-associated activating receptor [10], was recently reported on neutrophils [11]. FcγRIIIa and FcγRIIIb share 96% sequence identity in their extracellular IgG-binding regions [10], complicating efforts to delineate their distinct roles.

FcγRI (CD64) is the only high-affinity IgG receptor in humans [12]. Like FcγRIIIa, it signals through an ITAM-bearing adaptor. Depending on the cell type on which it is expressed, FcγRI effector responses include bacterial clearance, inflammation, anaphylaxis, endocytosis, phagocytosis, antigen presentation, release of B-cell stimulating factors, and anti-tumor responses (reviewed in [13]). Due to its high-affinity, FcγRI is likely saturated with circulating IgGs, which act like a threshold for receptor activation. However, it is likely that monomeric IgGs can be displaced by multivalent IgG immune complexes, causing receptor clustering which initiates downstream signaling [13].

In resting neutrophils, FcγRI is largely absent from the cell surface and stored in intracellular compartments [14, 15]. Its surface expression can be gradually induced by cytokines such as IFN-γ, G-CSF and GM-CSF [16–18], and increased surface FcγRI has also been observed during infections, where it is used clinically as a sensitive marker of sepsis [19–22]. More recently, we have shown that FcγRI can be rapidly mobilized to the neutrophil’s surface in response to immune complexes, enabling robust ROS production [15]. Yet, it remains unclear how FcγRI integrates with other FcγRs to shape context-specific effector responses, or whether its up-regulation supports normal immune function or drives pathogenic inflammation in chronic autoimmune disease. These gaps limit our understanding of neutrophil FcγRI biology.

Here, we sought to better define the role of FcγRI in neutrophil biology and inflammation. Specifically, we aimed to: (i) delineate FcγRI expression across circulating leukocyte populations; (ii) determine how neutrophil FcγRI surface expression relates to ROS production; and (iii) investigate whether FcγRI-mediated processes differ between healthy individuals and lupus patients, a population in which monocyte FcγRI is elevated [23]. Our work addresses gaps in understanding how FcγRI links immune complex recognition to neutrophil effector functions and evaluates this process in the context of health and disease.

## MATERIALS AND METHODS

*Materials.* Dextran-500, luminol, albumin from human serum (HSA), bovine serum albumin (BSA), cytochalasin B from *Drechslera dematioidea,* diphenylene iodonium (DPI) and Adenosine deaminase (ADA) were purchased from MilliporeSigma (Oakville, ON, Canada). Lymphocyte separation medium was purchased from Wisent (St-Bruno, QC, Canada). Lyophilized human immunoglobulins G (IgGs), purified from human plasma or serum by fractionation (purity of greater than 97% as determined by SDS-PAGE analysis) were purchased from Innovative Research, Inc (Novi, MI, USA). Pharm Lyse™ lysing solution was purchased from BD Biosciences (San Diego CA, USA). MitoTEMPO was purchased from Cayman Chemical (Ann Arbor, MI, USA). pHrodo Green *E. coli* BioParticles conjugates (P35366), lyophilized, were from Life Technologies Corporation (Eugene, OR, USA).

*Antibodies.* PercP-Cy5.5™-labeled mouse anti-human FcγRIII (3G8), PE-labeled mouse anti-human FcγRII (Clone FLI8.26, also known as 8.26), purified mouse anti-human FcγRI (10.1), V450-labeled mouse anti-human FcγRI (10.1), were purchased from BD Biosciences (San Jose, CA, USA). Monoclonal anti-human FcγRIIa (IV.3) was obtained from Bio X Cell (West Lebanon, NH, USA). This antibody recognizes a native extracellular epitope of FcγRIIa [24]. Anti-human albumin antibody produced in rabbit was purchased from Sigma-Aldrich (Oakville, ON, Canada).

*Ethics.* All experiments involving human tissues received approval from the research ethics committee of CHU de Québec-Université Laval (2022-6235).

*Preparation of heat-aggregated (HA)-IgGs.* HA-IgGs were freshly prepared each day as previously described [25], with modifications. Briefly, IgGs were resuspended in Hank’s Balanced Salt Solution (HBSS; 10 mM HEPES pH 7.4, 1.6 mM Ca^2+^ and no Mg^2+^), at a concentration of 25 mg/mL and heated at 63°C for 75 min to generate aggregates. The impact of Mg^2+^ was assessed in incubations and had no discernable effect on ROS production (**Suppl. Fig. S1**). Size assessment of formed HA-IgGS was performed by flow cytometry (**Suppl. Fig. S2**).

*Blood preparation for surface marker analysis.* No later than two hours after collection, venous blood was centrifuged at 280 × g for 10 min to remove platelet-rich plasma, then treated with HA-IgGs (1 mg/mL or indicated concentrations) for 15 min at 37°C. Following stimulation, 100 μL of blood was incubated with 1.9 mL of Pharm Lyse™ Buffer for 20 min at room temperature (RT) in the dark. After centrifugation (200 × g, 5 min), white blood cells were washed with 2 mL of phosphate-buffered saline (PBS).

*Flow cytometry.* Following appropriate treatments, cell pellets were resuspended with 100 µL of HBSS, and incubated with V450-labeled mouse anti-human CD64 (10.1), PE-labeled mouse anti-human CD32 (clone FLI8.26, also known as 8.26) and PercP-Cy5.5™-labeled mouse anti-human CD16 (3G8) for 30 min at 4°C. Cells were spun and resuspended in 400 µL of 1% paraformaldehyde before flow cytometry analysis. Acquisition was performed with a FACS Canto II flow cytometer with FACSDiva software, version 6.1.3 (BD Biosciences). For experiments with blood, monocytes, lymphocytes, and neutrophils were identified by forward and side scatter (FSC/SSC) gating. Where mentioned, samples were analyzed on a Cytek Northern Lights 3 lasers flow cytometer (V-B-R), with Spectroflo software, version 3.3.

*Isolation of human neutrophils.* Informed consent was obtained in writing from all donors. Data collection and analyses were performed anonymously. Neutrophils were isolated as originally described [26] with modifications [27]. Briefly, venous blood collected on isocitrate anticoagulant solution from healthy volunteers was centrifuged (250 x *g*, 10 min), and the resulting platelet-rich plasma was discarded. Leukocytes were obtained following sedimentation of erythrocytes in 2% Dextran-500. Neutrophils were then separated from other leukocytes by centrifugation on a 10 mL lymphocyte separation medium. Contaminating erythrocytes were removed using 20 sec of hypotonic lysis. Purified granulocytes (> 95% neutrophils, < 5% eosinophils) contained less than 0.2% monocytes, as determined by esterase staining. Viability was greater than 98%, as determined by trypan blue dye exclusion. The entire cell isolation procedure was carried out under sterile conditions at RT.

*Cell incubations.* Purified neutrophils were resuspended at a concentration of 10 x 10^6^ cells/mL at 37°C in HBSS supplemented with 10% human serum and 0.1 U/mL ADA to prevent the accumulation of endogenous adenosine in the medium, thus minimizing the previously demonstrated modulating effects of adenosine on inflammatory factor production by neutrophils [28–30].

*Preparation of IIC.* Immobilized immune complexes (IIC) were essentially prepared as described [31]. Briefly, high binding 96-well plates (Corning, NY, USA) were coated overnight at 4°C with 20 µg/mL HSA diluted in 50 mM carbonate/bicarbonate buffer (pH 9.6). Wells were washed with PBS containing 0.05% Tween 20 and subsequently blocked with PBS supplemented with 10% BSA for one hour at RT. Following blocking, wells were incubated for another hour at RT with rabbit anti-HSA antibodies diluted 1:400 in PBS. Finally, IIC-coated wells were washed twice with PBS containing 0.05% Tween 20 and once with HBSS prior to use.

*ROS detection.* ROS production was measured as previously described [32], with modifications. Briefly, neutrophils were resuspended at 1 × 10⁶ cells/mL in HBSS (10% human serum; 0.1 U/mL ADA). Luminol (10 µM final concentration) was added to neutrophil suspensions, and cells were treated as indicated. Cell suspensions (200 µL) were transferred into 96-well plates for exposure to HA-IgGs or IICs. For blood ROS experiments, 96-well white BRANDplates® (Brand GmbH + Co KG, Wertheim, BW, Germany) were used, with luminol at 1 mM. Plates were incubated at 37°C in the microplate reader Infinite M1000 PRO with i-control 2.0 software (Tecan, Morrisville, NC, USA). Luminescence intensity was monitored every 5 min for indicated times and the area under the curve (AUC) was determined. For experiments involving FcγR blockade, neutrophils were preincubated with anti-FcγRI (3 µg/mL) or anti-FcγRIIa (2.5 µg/mL) blocking antibodies for 10 min prior to stimulation with immune complexes. In specific instances, neutrophils were incubated with 125 µM cytochrome C and stimulated as indicated.

Where indicated, samples were preincubated for one hour with DPI (5 µM), or MitoTEMPO ((2-(2,2,6,6-Tetramethylpiperidin-1-oxyl-4-ylamino)-2-oxoethyl) triphenylphosphonium chloride, 100 µM).

*SLE cohort.* All samples and clinical data from patients used in this study were obtained from the biobank and database of systemic autoimmune rheumatic diseases (SARD) at CHU de Québec-Université Laval. Participants were diagnosed with SLE according to the 1997 Updated American College of Rheumatology (ACR) Classification Criteria [33, 34] and provided consent as required. Biological samples were paired with demographics and clinical data collected on the corresponding dates.

*Clinical and laboratory variables.* Demographic data, including age, biological sex and self-identified ethnicity, as well as lupus disease characteristics and medication use, were collected. Disease duration, organ damage (assessed using the Systemic Lupus International Collaborative Clinics/American College of Rheumatology [SLICC/ACR] Damage Index [35]), disease activity (assessed by the SLE Disease Activity Index – 2000 [SLEDAI-2K] [36]), and overall disease severity (assessed using the Lupus Severity Index [37] were recorded. Lupus nephritis was defined as the presence of the criteria for nephritis in the 2012 SLICC SLE classification criteria [38] and the report of lupus nephritis, classes I to VI, on a kidney biopsy report.

*Statistical analysis.* Statistical analysis was performed using GraphPad PRISM version 10 (GraphPad Software, San Diego, CA, USA). Where applicable, data are expressed as median ± interquartile range. Data were analyzed using tests including two-way ANOVA followed by Sidak’s multiple comparisons test, Wilcoxon signed-rank test, one-way ANOVA followed by Dunnett’s multiple comparisons test, Friedman test followed by Dunn’s multiple comparisons test, and Spearman’s correlation test, as indicated in the figure legends. For all analyses, differences were considered significant when *P* < 0.05.

## RESULTS

### Up-regulation of blood leukocyte FcγRI in response to immune complexes

To gain a comprehensive view of FcγR expression and modulation across leukocytes in response to immune complexes in circulation, we stimulated peripheral blood from healthy donors with HA-IgGs, used as a model of immune complexes [15], and assessed FcγRI, FcγRII, and FcγRIII expression by flow cytometry. In unstimulated blood, FcγRI was readily detectable on monocytes, modestly expressed on neutrophils, and marginally on lymphocytes (**Fig. 1A, pale blue**). Expression of FcγRII was strong on both neutrophils and monocytes, whereas FcγRIII expression was strongest on neutrophils. Following HA-IgG stimulation (1 mg/mL; 15 min; 37 °C), FcγRI up-regulation occurred predominantly on neutrophils, with a ∼3-fold increase in mean fluorescence intensity (MFI) and a 2.3-fold increase in the proportion of positive cells (**Fig. 1B, C, dark blue**). Monocytes also up-regulated their FcγRI, though to a lesser extent. FcγRII signal intensity levels decreased on both neutrophils and monocytes, whereas FcγRIII expression remained largely unchanged. Lymphocytes exhibited only marginal expression of all three receptors under the conditions tested.

**Figure 1.**
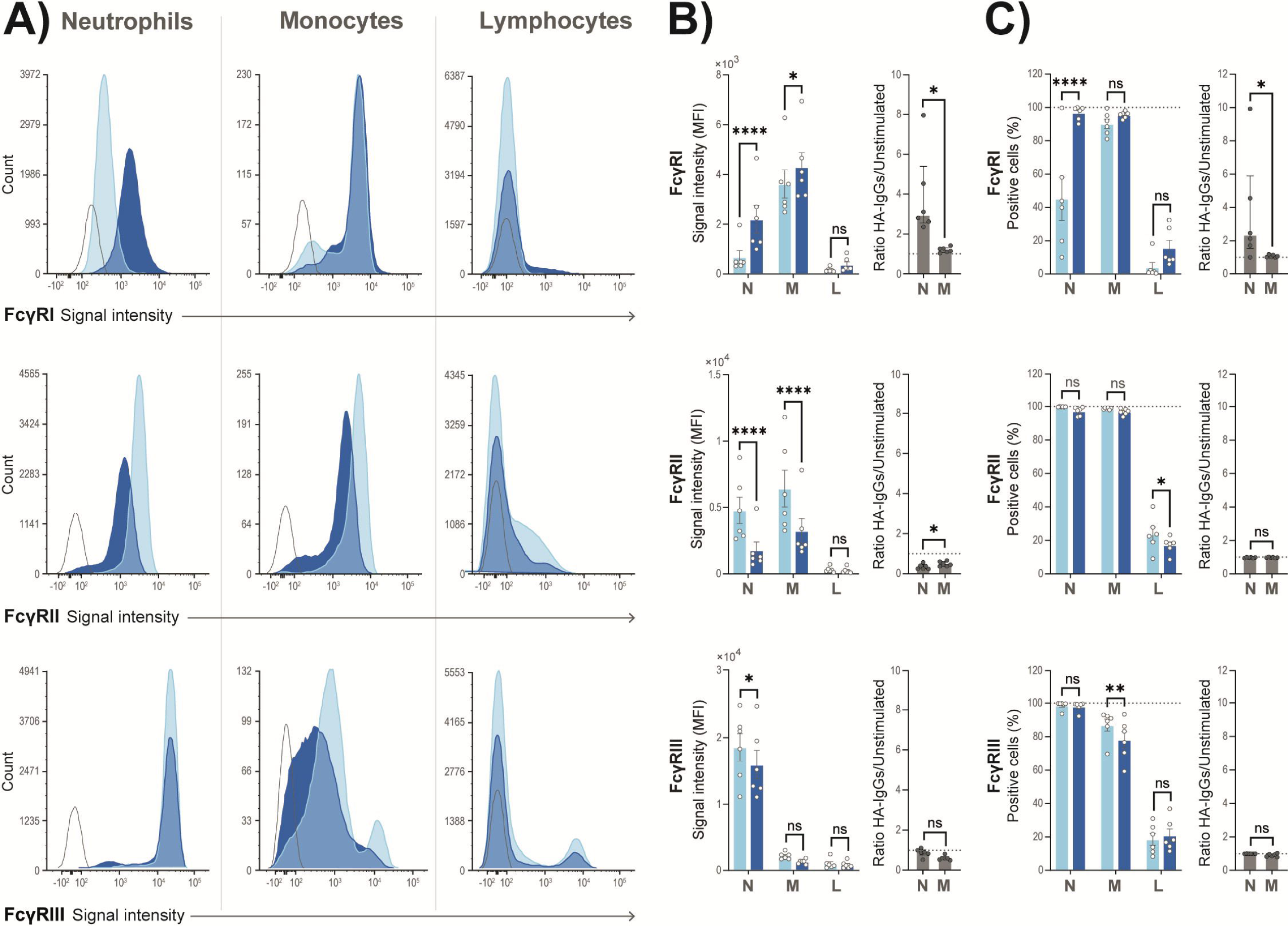
FcγRI is up-regulated distinctively on neutrophils upon incubation with HA-IgGs. Blood from healthy volunteers was left unstimulated (light blue) or stimulated with HA-IgGs (dark blue), and the surface expression of FcγRI, FcγRII, FcγRIII was measured on gated populations by flow cytometry, as described in *Methods.* **A)** Representative histograms from three leukocyte populations. Open histogram: negative control. **(B–C)** Data shown as individual values from six different donors, with median and interquartile range for mean fluorescence intensity (MFI, **B**) and percentage of positive cells (**C**). Statistical analysis was performed using the two-way ANOVA followed by Sidak’s multiple comparisons test (signal intensity graphs) or the Wilcoxon test (ratio graphs) (ns: not significant, \**P* < .05, \*\*\*\**P* < .0001). N: Neutrophils, M: Monocytes, L: Lymphocytes, HA-IgGs: Heat-Aggregated IgGs.

### FcγRI contributes to immune complex–induced ROS generation in neutrophils

Next, we investigate the functional consequences of neutrophil FcγRI up-regulation on immune complex–induced ROS production using neutrophil-enriched cell suspensions. Previous work by Behnen *et al.* reported FcγRI-independent ROS generation in neutrophils exposed to immobilized immune complexes (IIC) [31]. In that study, the authors pre-incubated neutrophils with an anti-FcγRI antibody and washed it away before exposing cells to IIC. When reproducing the key washing step prior to stimulation (i.e., washing antibodies off prior to incubation with HA-IgGs), we obtained similar results: ROS production was not significantly altered (**Fig. 2A**). However, when leaving the blocking anti-FcγRI antibody present during stimulations, ROS production was significantly reduced, returning close to non-stimulated levels. A similar pattern was observed when blocking FcγRI on neutrophils stimulated with IIC (**Fig. 2B**). As expected [5, 31], blocking FcγRIIa abolished ROS production regardless of whether the antibody was washed away or left in (**Fig. 2C**).

**Figure 2.**
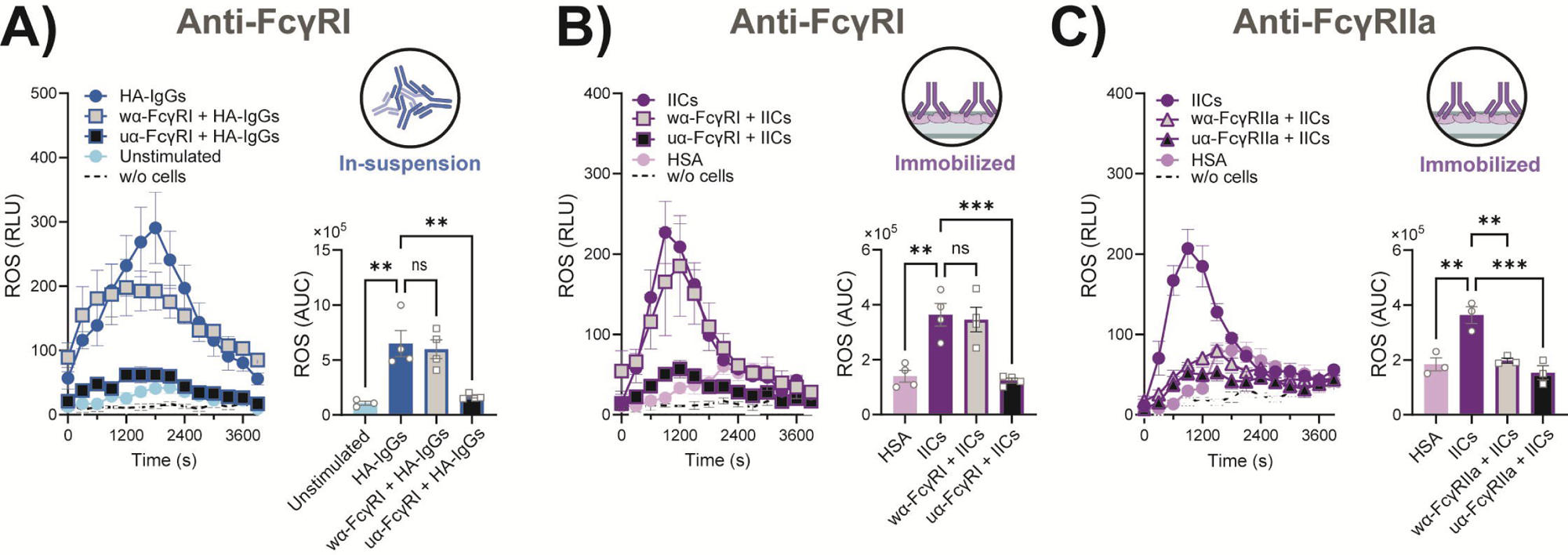
ROS production by human neutrophils exposed to immune complexes is dependent on FcγRI. Neutrophil-enriched cell suspensions were pre-incubated for 10 min with anti-FcγRI (**A, B**) or anti-FcγRIIa (**C**) blocking antibodies, which were either washed away (wα-FcγRI, wα-FcγRIIa) before exposure to immune complexes, or left present during stimulation (unwashed, uα-FcγRI, uα-FcγRIIa) with HA-IgGs (blue) or immobilized immune complexes (purple). In each panel, line graphs show ROS kinetics (relative luminescence units, RLU) with median and interquartile range, while bar graphs show the area under the curve (AUC) with individual values, median, and interquartile range. Data are from at least three different donors. Statistical analysis was performed using one-way ANOVA followed by Dunnett’s multiple comparisons test (ns: not significant, \*\**P* < .01, \*\*\**P* < .001). HA-IgGs: Heat-Aggregated IgGs, HSA: Human Serum Albumin, IICs: Immobilized Immune Complexes, ROS: Reactive Oxygen Species. Created in BioRender. Huot, S. (2025) https://BioRender.com/sdpy77q (Licence number: NP28SWS6IY).

To clarify whether the magnitude of ROS increase observed with our HA-IgG model is biologically meaningful, a series of experiments was performed using lyophilized, opsonized *E. coli* prestained with the pHrodo probe, whose fluorescence increases in acidic conditions, such as within phagolysosomes. Upon incubation with neutrophils, opsonized particles stimulated ROS production in a concentration-dependent manner (**Suppl. Fig. S3A**). This incubation resulted in bacteria uptake and internalization, and in the up-regulation of FcγRI (**Suppl. Fig. S3B, C**). In some cases, lyophilized, opsonized *E. coli* were stained with propidium iodine (PI), prior to incubation with neutrophils, to assess bacterial DNA integrity. As can be seen in **supplemental figure S3D**, PI signal further increased when ROS generation was blocked with DPI, indicating the inhibition of bacterial destruction. As observed with HA-IgGs, opsonized bacteria caused the up-regulation of FcγRI (**Suppl. Fig. S3C**) and ROS generation (**Suppl. Fig. S3E**). Its blockade dramatically prevented both bacterial uptake and ROS generation (**Suppl. Fig. S3B, E**). Comparison between cytochrome C reduction and Luminol-based assays yielded similar patterns (**Suppl. Fig. S4**) and indicated that Luminol-based ROS species detected are from intracellular sources **(Suppl. Fig. S5)**.

### Degranulation enhances FcγRI surface mobilization and amplifies ROS production

Inhibition of exocytosis prevents neutrophil FcγRI up-regulation and ROS production [15], supporting a functional link between FcγRI and this cell response. Here, we tested the impact of enhanced degranulation on neutrophil FcγRI expression and ROS production with cytochalasin B (CB), a known degranulation enhancer acting through actin disruption [39]. On its own, CB (10 µM) had no effect on FcγR expression. However, in the presence of HA-IgGs, CB potentiated both FcγRI up-regulation and ROS production in a concentration-dependent manner and concomitantly reduced FcγRIII (**Fig. 3A–C**). Whereas FcγRI up-regulation and FcγRIII downregulation were significant for both signal intensity and percentage of positive cells, FcγRII changes only reached significance for the percentage of positive cells.

**Figure 3.**
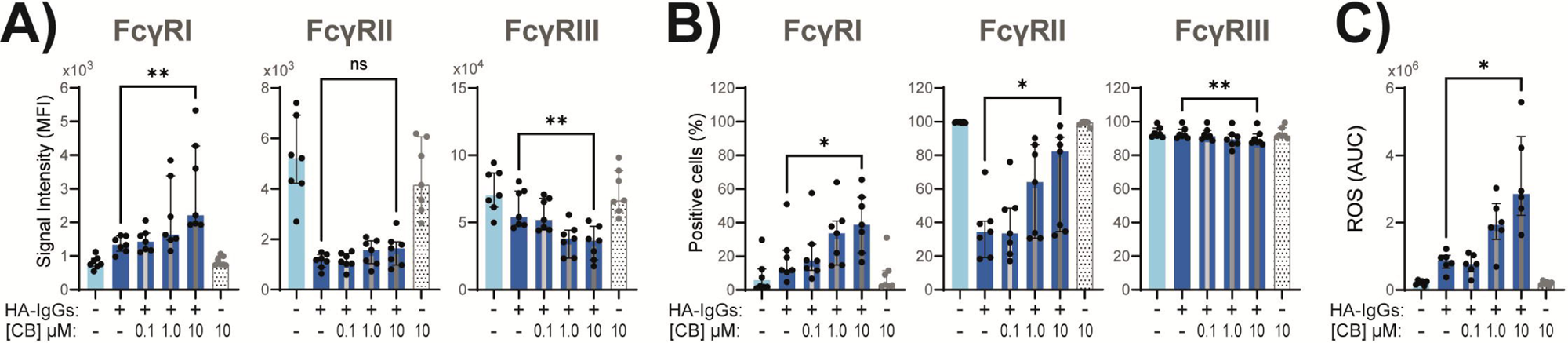
Cytochalasin B potentiates FcγRI surface mobilization and ROS production in human neutrophils. Neutrophil-enriched cell suspensions were left unstimulated or pre-incubated with the indicated concentrations of cytochalasin B (CB), with or without stimulation by heat-aggregated IgGs (HA). FcγR surface expression was measured by flow cytometry as signal intensity (**A**) and percentage of positive cells (**B**), and ROS production was assessed by luminol-based chemiluminescence (**C**). Data are shown as individual values from at least six different donors, with median and interquartile range. Statistical analysis was performed using the Friedman test followed by Dunn’s multiple comparisons test (ns: not significant, \**P* < .05, \*\**P* < .01). AUC: Area Under the Curve; MFI: Mean Fluorescence Intensity; ROS: Reactive Oxygen Species.

Collectively, these findings support a model in which initial engagement of low-affinity FcγRIIa triggers FcγRI mobilization from intracellular secretory compartments. This mobilization leads the to up-regulation of FcγRI on the neutrophil surface, which in turn contributes to ROS generation in response to immune complexes. While the Syk- and PI3K pathways may be involved [14, 15], further studies are needed to fully dissect the signaling pathway(s) regulating FcγRI surface expression, in neutrophils. To determine whether generation of ROS contribute to FcγRI surface up-regulation, neutrophils were treated with DPI, a competitive inhibitor of flavin-dependent enzymes that blocks NADPH oxidase (NOX) activity [40]. While DPI markedly inhibited ROS production, mitoTEMPO, a mitochondria-targeted superoxide dismutase mimetic [41], had little effect, confirming that the detected ROS are primarily of NADPH oxidase origin rather than mitochondrial (**Fig. 4A**). Importantly, inhibition of ROS by DPI did not impair FcγRI surface mobilization, indicating that FcγRI up-regulation occurs independently of ROS production (**Fig. 4B**). No significant impact of DPI on FcγRII was observed either (**Fig. 4C**).

**Figure 4.**
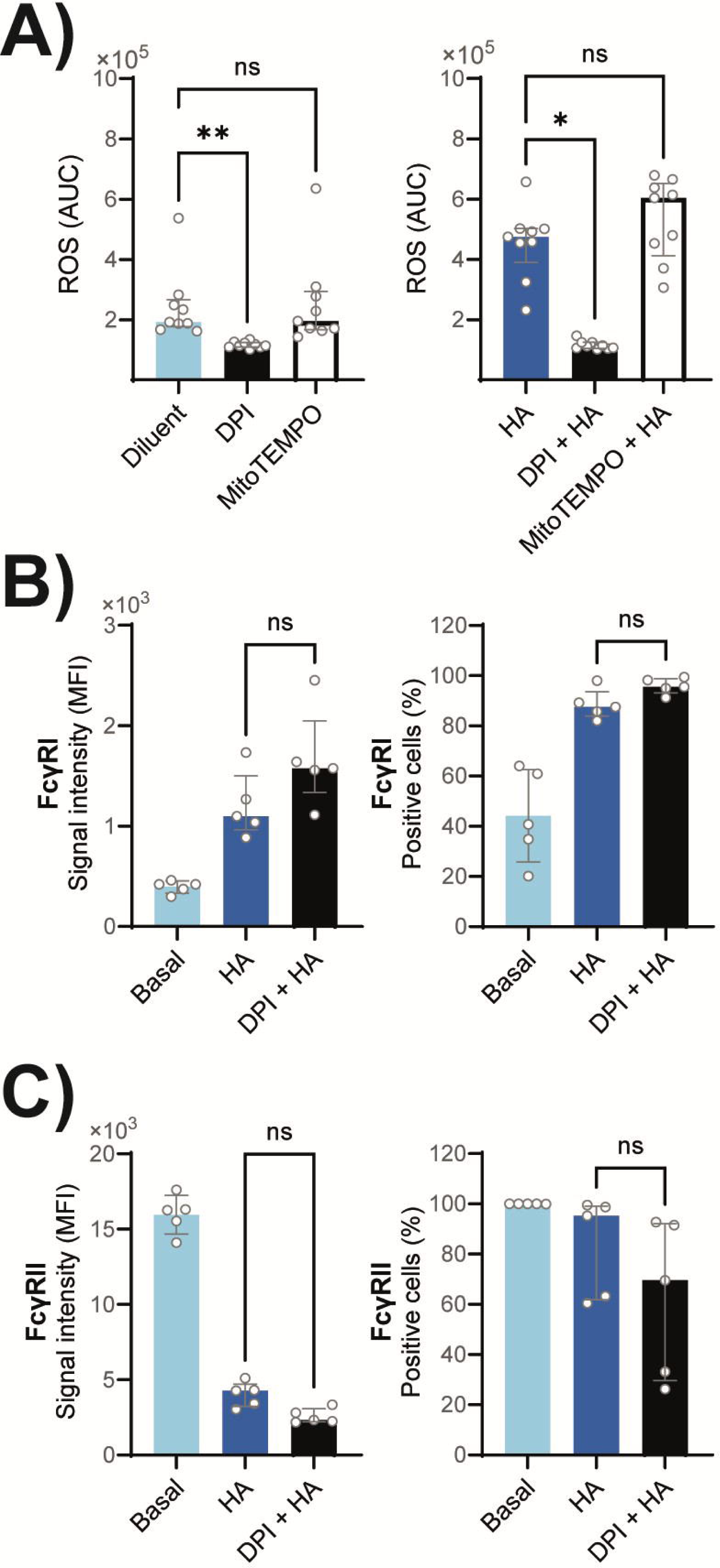
**DPI inhibits ROS production without affecting FcγRI up-regulation in human neutrophils**. Neutrophil-enriched cell suspensions were pre-incubated for 1 h with DPI, mitoTEMPO, or diluent, then left unstimulated (basal) or stimulated with HA-IgGs (HA). **A**) ROS production was assessed by luminol-based chemiluminescence, as described in *Methods.* **B, C**) FcγR surface expression was assessed by flow cytometry for mean fluorescence intensity (MFI) and percentage of positive cells. Data are shown as individual values from at least five different donors, with median and interquartile range. Statistical analysis was performed using the Friedman test followed by Dunn’s multiple comparisons test (ns: not significant, \**P* < .05, \*\**P* < .01). DPI: diphenylene iodonium, HA-IgGs: Heat-Aggregated IgGs, ROS: Reactive Oxygen Species, w/o: without.

### Neutrophil FcγRI expression correlates with ROS production in health and disease

So far, our studies linking neutrophil FcγRI and ROS generation were primarily based on high concentrations of aggregated IgGs. To determine whether this would also occur in a more physiologic setting, we first stimulated blood samples with HA-IgG concentrations comparable to those reported for immune complexes in circulation [42]. Physiologically relevant HA-IgG concentrations were sufficient to significantly increase ROS production and FcγRI surface expression, in a concentration-dependent manner (**Fig. 5**).

**Figure 5.**
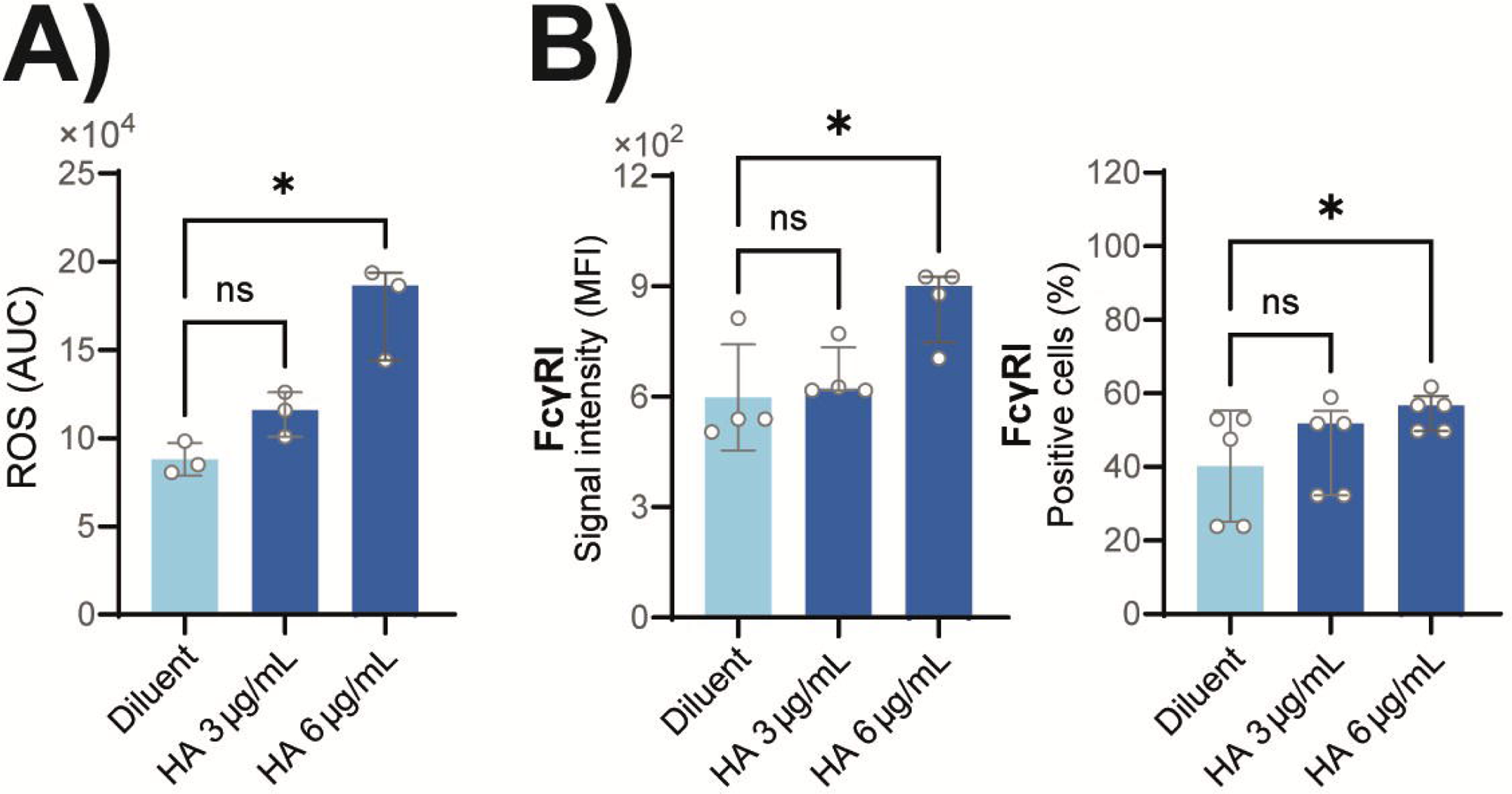
Physiologically relevant HA-IgG levels enhance neutrophil FcγRI expression and ROS generation in blood. Fresh blood samples from healthy donors were incubated with diluent or with indicated concentrations of HA-IgGs (HA). ROS production was assessed by luminol-based chemiluminescence (**A**) and neutrophil-FcγRI surface expression was measured by flow cytometry for signal intensity and percentage if positive cells (**B**). Data are shown as individual values from at least three different donors, with median and interquartile range. Statistical analysis was performed using the Friedman test followed by Dunn’s multiple comparisons test (ns: not significant, \**P* < .05). AUC: Area Under the Curve, HA-IgGs: Heat-Aggregated IgGs, MFI: Mean Fluorescence Intensity, ROS: Reactive Oxygen Species.

Given the substantial evidence presented, suggesting a functional link between neutrophil FcγRI and ROS, we measured FcγRI surface expression and basal ROS generation in whole blood samples from 12 healthy volunteers. ROS levels significantly correlated with neutrophil FcγRI expression, as measured by MFI and by percentage of FcγRI-positive cells (**Fig. 6A**). Neutrophil FcγRII and FcγRIII expression showed no correlation, and none of the monocyte Fcγ receptors (FcγRI, FcγRII, or FcγRIII) either (**Fig. 6B**).

**Figure 6.**
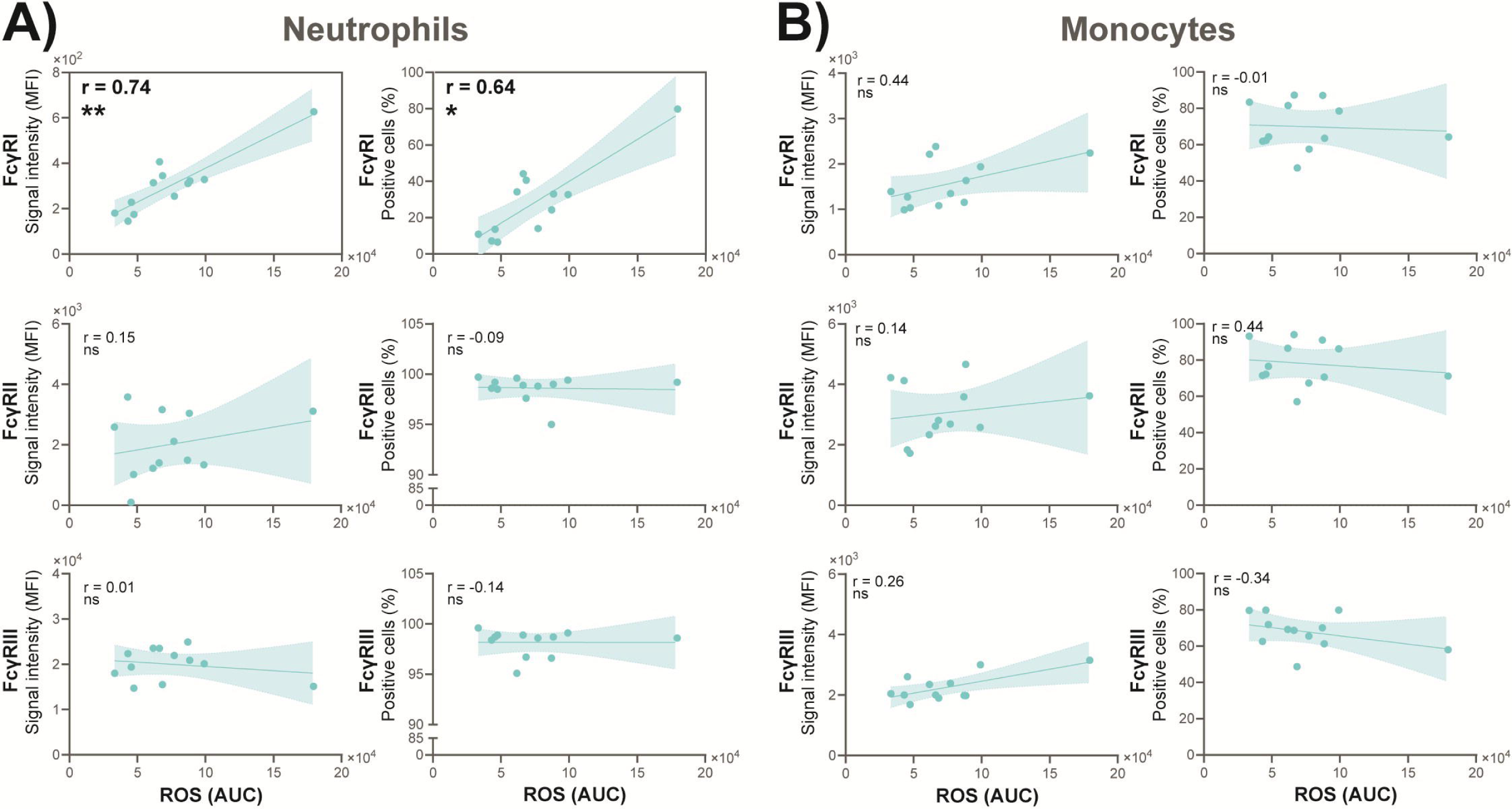
Neutrophil FcγRI surface expression correlates with ROS production in healthy donor blood. Fresh blood samples from healthy donors (n = 12) were assessed for ROS production by luminol-based chemiluminescence and for FcγR surface expression on flow cytometry–gated neutrophils (**A**) and monocytes (**B**). Correlations were evaluated using Spearman’s rank test (r values and *P* indicated; ns: not significant, \**P* < .05, \*\**P* < .01). Linear regression lines with 95% confidence intervals are shown for visualization. AUC: Area Under the Curve, MFI: Mean Fluorescence Intensity, ROS: Reactive Oxygen Species.

Monocyte FcγRI expression is a biomarker of disease activity in systemic lupus erythematosus [23, 43]; neutrophil FcγRI has received less attention in this context. In turn, we measured the association between myeloid FcγRI and ROS generation in fresh blood samples from 12 age- and sex-matched patients with systemic lupus erythematosus (lupus). Characteristics of this group of patients are presented in **Table 1** **and supplemental fig. S6**). Circulating neutrophil FcγRI expression also correlated with ROS levels in patients with lupus, as assessed by both MFI and percentage of FcγRI-positive cells (**Fig. 7A**). Monocytes showed a weaker pattern, but significance was reached for MFI (**Fig. 7B**). Together, these findings identify neutrophil FcγRI expression as a key determinant of oxidative response, both in healthy volunteers and in patients with lupus.

**Figure 7.**
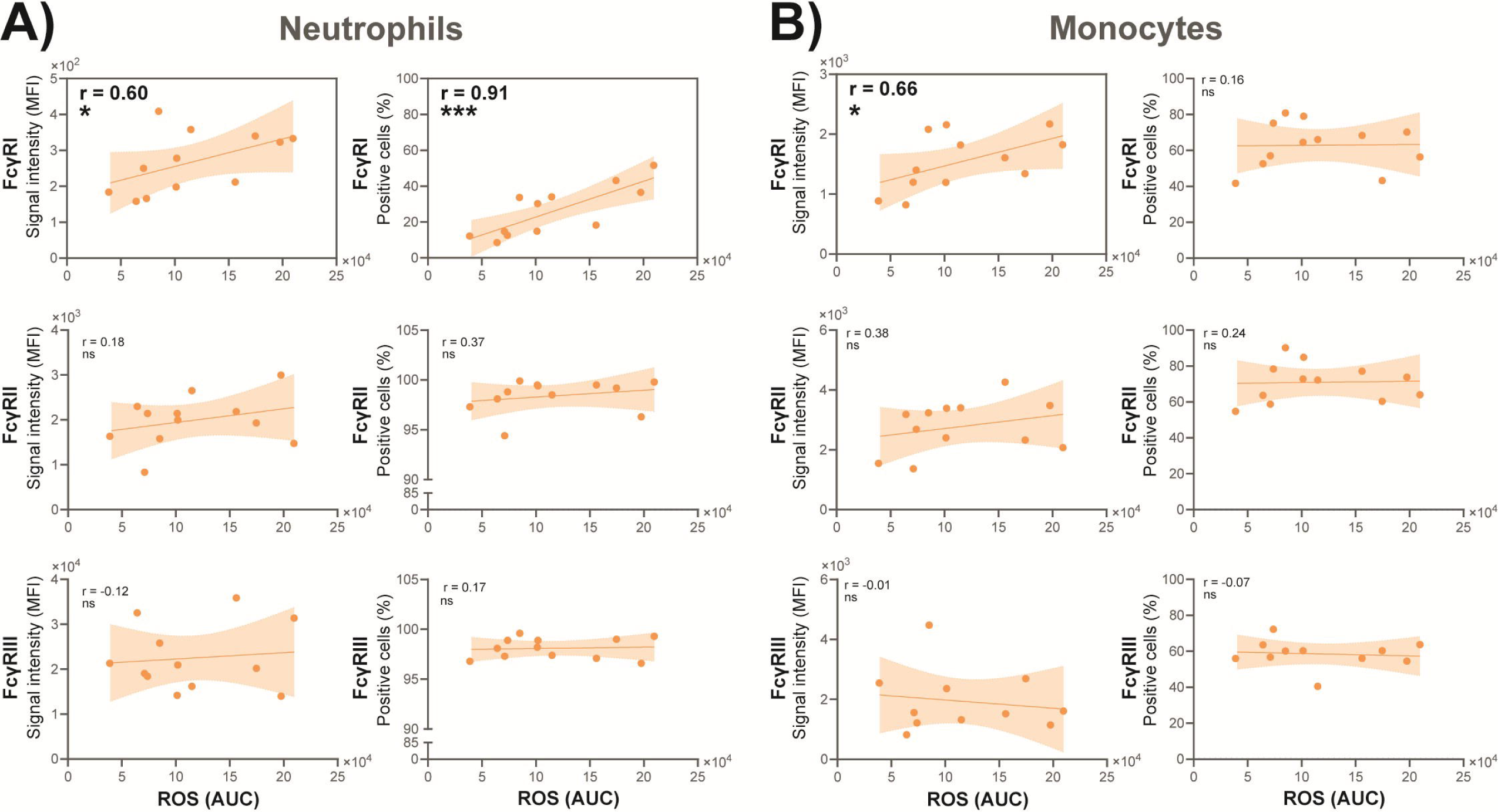
Neutrophil FcγRI surface expression correlates with ROS production in blood from patients with systemic lupus erythematosus. Fresh blood samples from patients with lupus (n = 12) were assessed for ROS production by luminol-based chemiluminescence and for FcγR surface expression on flow cytometry–gated neutrophils (**A**) and monocytes (**B**). Correlations were evaluated using Spearman’s rank test (r values and *P* indicated; ns: not significant, \**P* < .05, \*\**P* < .01, \*\*\**P* < .001). Linear regression lines with 95% confidence intervals are shown for visualization. AUC: Area Under the Curve; MFI: Mean Fluorescence Intensity; ROS: Reactive Oxygen Species.

**Table 1.**
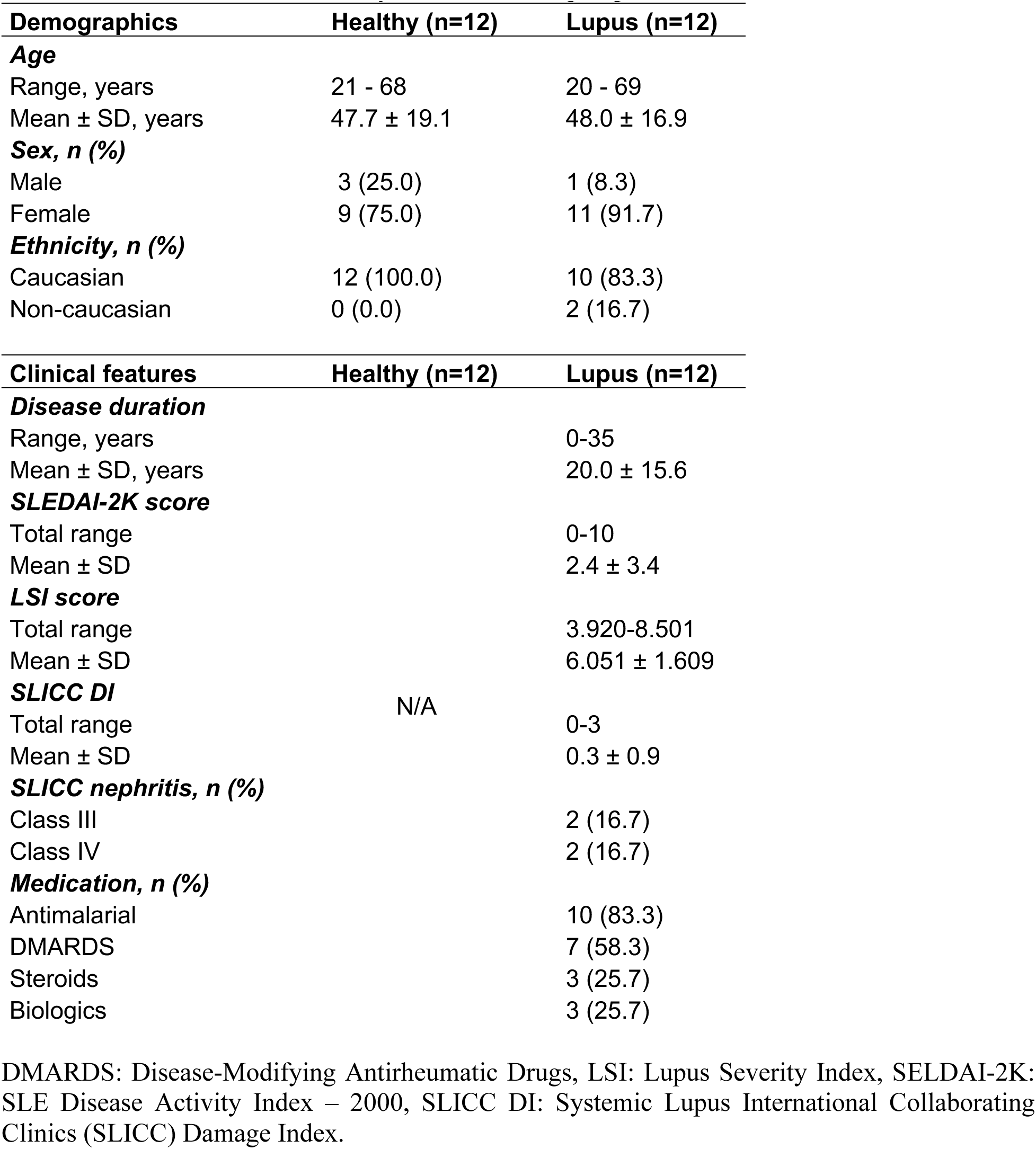
Characteristics of healthy controls and lupus patients

## DISCUSSION

Previous work from our group demonstrated that HA-IgGs can mobilize FcγRI to the surface of isolated neutrophils [15]. Here, we extend these findings by showing that FcγRI is the most dynamically regulated Fcγ receptor on neutrophils, compared to monocytes and lymphocytes in circulation, in response to HA-IgGs. Its surface expression correlates with measurable intracellular ROS generation. Contrasting with the slower, cytokine-driven up-regulation previously described [16–18], this rapid response documents a new regulation of acute neutrophil activation.

ROS production is a hallmark neutrophil response to immune complexes, and is chiefly attributed to FcγRIIa [9, 36]. The evidence presented in this study suggests that FcγRI also regulates this function. On the one hand, enhanced exocytosis amplified both FcγRI surface expression and ROS production, in line with our previous work showing that degranulation is required for FcγRI surface mobilization [15], and highlights FcγRI’s capacity to be quickly deployed. On the other hand, blocking either FcγRI or FcγRIIa during neutrophil stimulation abolished oxidative responses. The sequential FcγRI up-regulation in response to FcγRIIa activation exemplifies how-FcγRs can cooperate to generate context-specific responses [44], with FcγRI acting as a key regulator of ROS production in neutrophils [4] rather than an initial sensor. ROS production was confirmed to originate mainly from cellular NADPH oxidase activity, underscoring the canonical oxidative burst machinery as the effector downstream of FcγRI. Our observation that FcγRII levels decreased while FcγRI increased following exposure to HA-IgGs suggests a dynamic redistribution of receptor usage, possibly shifting neutrophils toward high-affinity FcγRI-driven responses under conditions of sustained immune complex stimulation. The expression of FcγRIII remained largely stable throughout incubations.

ROS production by neutrophils was demonstrated to be dependent, in part, on FcγRI both in response to aggregates in suspension and to immobilized immune complexes. This indicates that FcγRI might be instrumental, not only on neutrophils in circulation, but also in those migrated and adherent to tissues. In support of this, FcγRI up-regulation has been observed on neutrophils in response to CD11b or CD18 ligation [45]. Furthermore, neutrophil FcγRI surface expression consistently reflected oxidative capacity, extending earlier findings that FcγRI-positive human neutrophils generate more ROS than their FcγRI-negative counterparts [46], positioning FcγRI as a central regulator of ROS production in neutrophils.

A dual role of FcγRI is apparent: on the one hand, its rapid mobilization likely provides an effective mechanism against IgG-opsonized particles. On the other hand, in autoimmune diseases where immune complexes are abundant or immobilized in tissues, FcγRI may drive pathological neutrophil activation and oxidative tissue injury. The balance between these outcomes may depend on the local cytokine milieu, the density and composition of immune complexes, and the interplay with other cells. Thus, our findings may have implications for diseases characterized by immune-complex deposition and systemic inflammation, such as lupus. As the only high affinity IgG receptor in humans, FcγRI is a target in many clinical settings, including neuropathy, cancer, and in autoimmune diseases [13]. In lupus, monocyte FcγRI is firmly associated with key clinical features and progression [23, 43, 47, 48]. Neutrophil activation is held to contribute to lupus pathogenesis through tissue damage, NET formation, and perpetuation of inflammation [49]. However, information regarding neutrophil FcγRI and lupus is less clear. In the present study, although monocyte FcγRI correlated with ROS production in patients living with lupus, correlations between FcγRI expression and oxidative responses were stronger in neutrophils, both in healthy subjects and in patients. Given the well-documented involvement of monocyte FcγRI in lupus, this finding was rather surprising. While no difference in FcγRI expression was observed between the two groups, the fact that neutrophils outnumber monocytes by a 10-to-1 factor in the circulation, and that most of the ROS produced in blood originates from neutrophils, suggest that neutrophil FcγRI may be a more immediate regulator of inflammatory effector functions, both in health and disease. Our work opens several avenues for further investigation. The regulation of FcγRI and ROS production by cytokines, complement components, or other inflammatory mediators *in vivo* warrants closer examination. It may be important to determine whether FcγRI-dependent ROS production contributes to NETosis, a process increasingly recognized in lupus pathogenesis [50].

Although useful for controlled stimulation, HA-IgGs do not fully recapitulate the structural diversity and antigen-specificity of, for example, opsonized microorganisms [51]. Also, additional mechanistic studies will be important to dissect how FcγRI activity integrates with that of other receptors such as FcγRIIa, FcγRIIIa, FcαR, and possibly others [5, 11, 52–54] to reveal better targeted therapeutic opportunities. Finally, the modest size of our cohorts prevents one from drawing population-wide conclusions or correlations with clinical parameters such as disease activity or organ involvement in the lupus group. Larger, longitudinal studies will be required to establish whether neutrophil FcγRI expression serves as a reliable biomarker for ROS generation. The possible contribution of low-density granulocytes (LDGs) has not been addressed. LDGs are implicated in the pathogenesis of lupus and other autoimmune diseases, for their association with vascular damage [55]. Specific markers to identify LDGs in circulation or in tissue will be useful to evaluate their impact on FcγRI biology [56]. From a translational standpoint, selective modulation of FcγRI could represent a therapeutic strategy: blocking its activity may limit pathological oxidative responses in autoimmune disease, while enhancing its mobilization could improve host defense against infections. Immunotoxins targeting FcγRI have already demonstrated significant clinical potential to resolve chronic inflammation [57–61]. Approaches to specifically skew neutrophil responses away from oxidative damage and toward resolution, may offer novel opportunities in the treatment of immune complex–mediated conditions [62].

In conclusion, we identify neutrophil FcγRI as an additional cellular component regulating the oxidative response machinery. Its strong association with ROS production in both health and disease support a central role for FcγRI in neutrophil biology and may represent both a biomarker and a therapeutic target in autoimmune disease. By uncovering this pathway, our findings provide a new perspective on how immune complex–driven inflammation is regulated by neutrophils.

## ACKNOWLEDGEMENTS

This work was funded by a grant from the Arthritis Society, Canada (21-0000000121) to MP, and by the Fonds de recherche du Québec (FRQ) through the research centre grant for the CHU de Québec-Université Laval Research Center (30641). SH is the recipient of a training award from the Fonds de recherche du Québec – Santé (FRQS; doi: 10.69777/268262), a CIHR-Frederick Banting and Charles Best Canada Graduate Scholarship, a Centre ARThrite-UL studentship, and a *Bourse de Formation Desjardins pour la recherche et l’innovation* (CHU de Québec). PRF holds a Tier 1 Canada Research Chair in Systemic Autoimmune Rheumatic Diseases. The authors thank Prof. Martin Pelletier for kindly providing DPI and *E. coli* particles.

## AUTHOR CONTRIBUTIONS

M. Pouliot, S. Huot, and P.R. Fortin conceived and designed the research. M. Pouliot, S. Huot and C. Laflamme designed the experiments and analyzed and interpreted the data. S. Huot and C. Laflamme performed the experiments. M. Pouliot and S. Huot were involved in drafting and writing the manuscript. All authors approved the final version of the manuscript.

Competing interests: The authors declare that they have no competing interests.

Data availability: Data sharing not applicable to this article as no datasets were generated or analyzed during the current study.

PREPRINT: This work is deposited here [63]: DOI: doi.org/10.1101/2025.10.01.679808

