## Supplemental materials for "Neutrophil FcγRI expression as a determinant of oxidative responses in human blood"

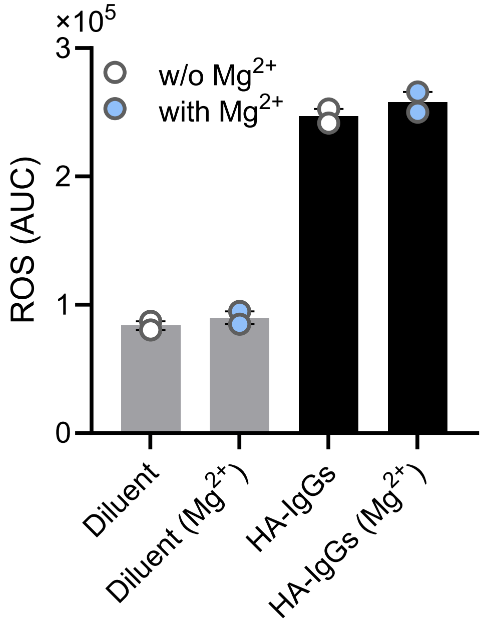


**Supplemental Figure S1. Absence of $Mg^{2+}$ during cell incubations has no discernible impact on neutrophil ROS production.** Neutrophil-enriched cell suspensions were incubated in HBSS supplemented with 10% human serum and 0.1 U/mL ADA, in the absence (w/o $Mg^{2+}$) or presence of 0.1 g/L $MgCl_2 \cdot 6H_2O$ (with $Mg^{2+}$) and stimulated with HA-IgGs. ROS production was assessed by luminol-based chemiluminescence as described in *Methods*. Data are shown as individual values from two different donors, with median and interquartile range. AUC: Area Under the Curve, HA-IgGs: Heat-Aggregated IgGs, $Mg^{2+}$: ($MgCl_2 \cdot 6H_2O$), 0.1 g/L, as formulated in Wisent Inc. HBSS, (Catalog no. 311-515), ROS: Reactive Oxygen Species,

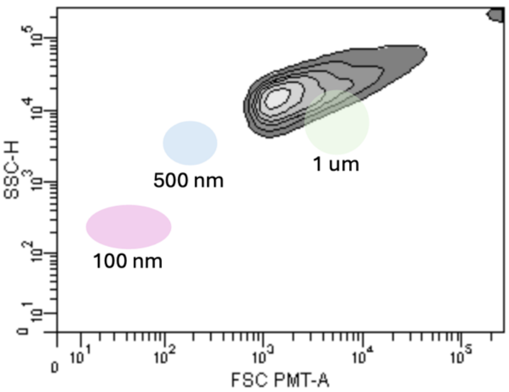


**Supplemental Figure S2. Size determination of the HA-IgG preparations**. HA-IgGs were prepared and analyzed by flow cytometry, as described in *Methods*. Results are displayed by side scatter (SSC)-H (granularity) and forward scatter (FSC)**.** Silica beads of various size (pink, 100 nm; blue, 500 nm, and green, 1 micron) were used as reference (G. Marcoux, A. Magron, C. Sut, A. Laroche, S. Laradi, H. Hamzeh-Cognasse, et al. Transfusion 2019 Vol. 59 Issue 7 Pages 2403-2414. PMID: 30973972). Results are from n=1 typical experiment. The vast majority of HA-IgGs formed (shades of gray) were just below 1 micron in diameter.

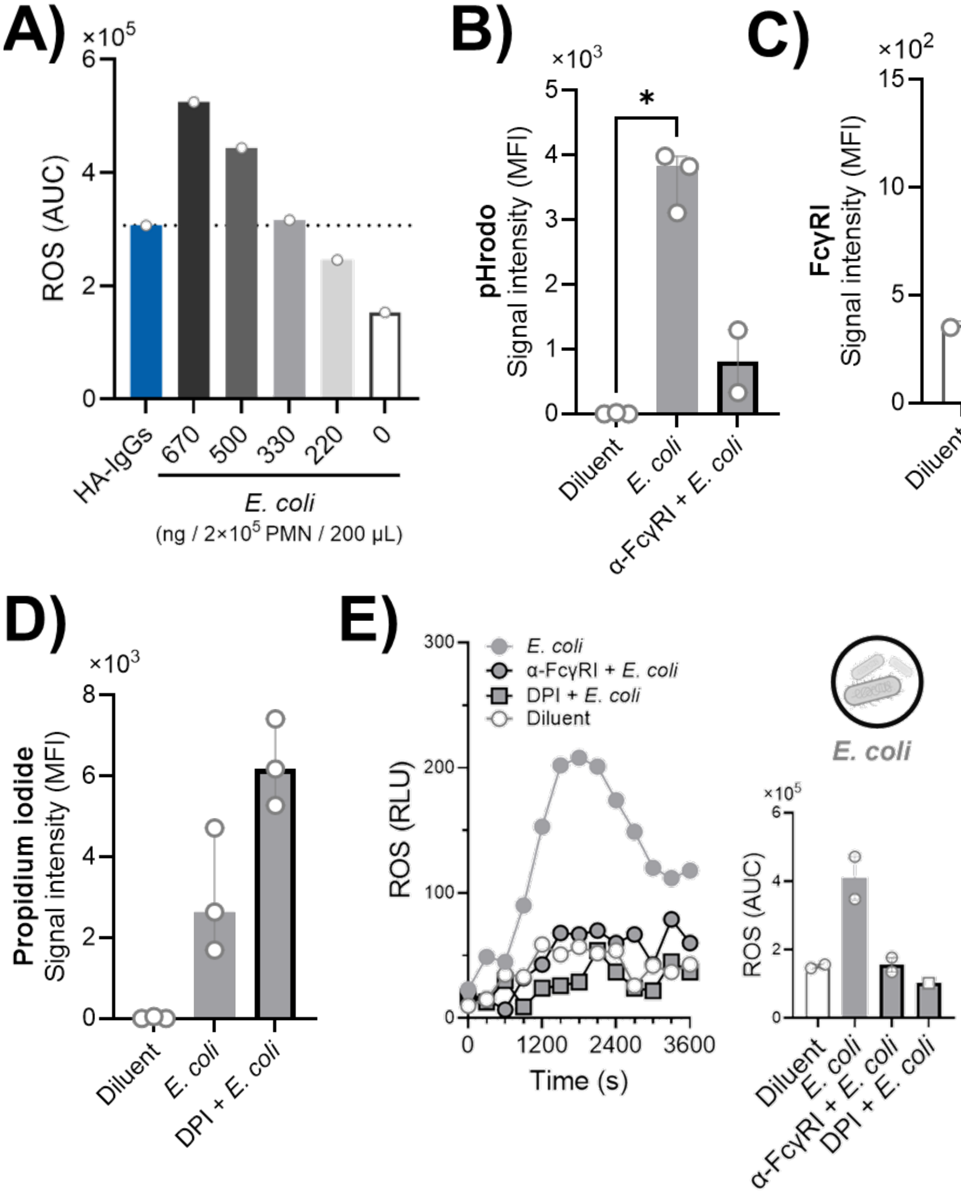

A)
×10^5
ROS (AUC)
HA-IgGs
670
500
330
220
0
E. coli
(ng / 2×10^5 PMN / 200 µL)
B)
×10^3
pHrodo
Signal intensity (MFI)
*
Diluent
E. coli
α-FcγRI + E. coli
C)
×10^2
FcγRI
Signal intensity (MFI)
Diluent
E. coli
DPI + E. coli
D)
×10^3
Propidium iodide
Signal intensity (MFI)
Diluent
E. coli
DPI + E. coli
E)
E. coli
α-FcγRI + E. coli
DPI + E. coli
Diluent
ROS (RLU)
Time (s)
E. coli
×10^5
ROS (AUC)
Diluent
E. coli
α-FcγRI + E. coli
DPI + E. coli

**Supplemental Figure S3. The amplitude of ROS production contributes to bacteria degradation. A)** Neutrophil-enriched cell suspensions were incubated with indicated concentrations of opsonized pHrodo green-stained *E. coli*, and ROS production was measured. **B, C)** Cells were incubated with opsonised bacteria (1.65 µg per 1 x $10^6$ PMNs; equivalent to 330 ng *E. coli* per 2 × $10^5$ cells) for 30 min at 37 °C, spun and resuspended in PBS. Cells were directly analysed for pHrodo signal by flow cytometry (B), or stained with V450-labeled mouse anti-human CD64 (FcγRI), as described in *Methods*, prior to flow cytometry acquisition (C). **D)** Opsonized bacteria were pre-stained with propidium iodide for 10 min prior to incubation with cells and analysis by flow cytometry. **E)** Cells were incubated with opsonised bacteria (1.65 µg per 1 x $10^6$ PMNs) before ROS analysis. Where indicated, samples were preincubated for 15 min with anti-FcγRI (3 µg/mL), or for one hour with DPI (5 µM). A, E: Data are shown as one representative experiment or as individual values from two independent donors with median and interquartile range. B,D: Flow cytometry data were acquired on a Cytek Northern Lights 3 lasers (V-B-R), with the Spectroflo software, version 3.3. B,C,D: Data are shown as individual values from at least two different donors with median and interquartile range. Statistical analysis was performed using the Kruskal–Wallis test ($P < 0.05$). AUC: Area Under the Curve, DPI: Diphenylene Iodonium, HA-IgGs: Heat-Aggregated IgGs, MFI: Mean Fluorescence Intensity, PBS: Phosphate-Buffered Saline PMN: polymorphonuclear neutrophil, RLU: Relative Luminescence Unit, ROS: Reactive Oxygen Species. pHrodo green-stained *E. coli* bacteria were kindly provided by Dr. Martin Pelletier.

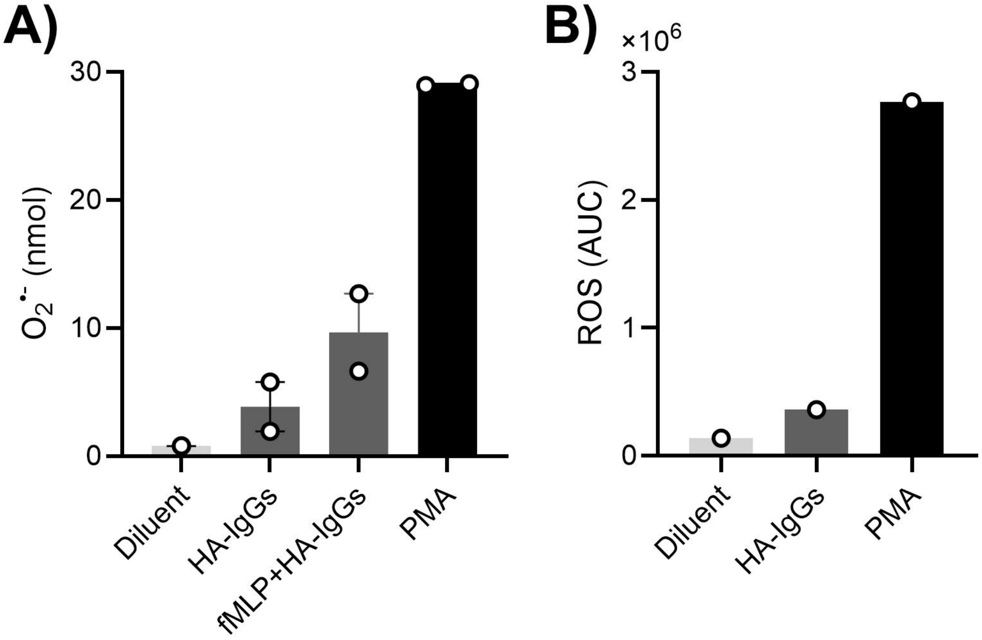


**Supplemental Figure S4. Cytochrome C reduction and luminol-enhanced chemiluminescence yield similar stimulus-dependant ROS production patterns by neutrophils. A)** Neutrophil-enriched cell suspensions were resuspended at 1 x $10^6$ cells/mL in HBSS supplemented with 125 µM cytochrome C. Cells were preincubated ± fMLP (100 nM) for 10 min at 37 °C before stimulation with the indicated components for an additional 10 min. Cells were lysed (0.1% Igepal), spun, and the supernatant transferred to a 96-well plate for optical density measurement (550 nm with correction at 540 nm). Superoxide anion production was calculated using the formula published by Dahlgren and Karlsson (C. Dahlgren and A. Karlsson. J Immunol Methods 1999 Vol. 232 Issue 1-2 Pages 3-14. PMID: 10618505). **B)** ROS production was assessed by luminol-based chemiluminescence as described in *Methods*. Data are shown as individual values from two different donors with median and interquartile range (panel A); panel B shows a single donor. AUC: Area Under the Curve, HA-IgGs: Heat-Aggregated IgGs, PMA: Phorbol 12-Myristate 13-Acetate, ROS: Reactive Oxygen Species

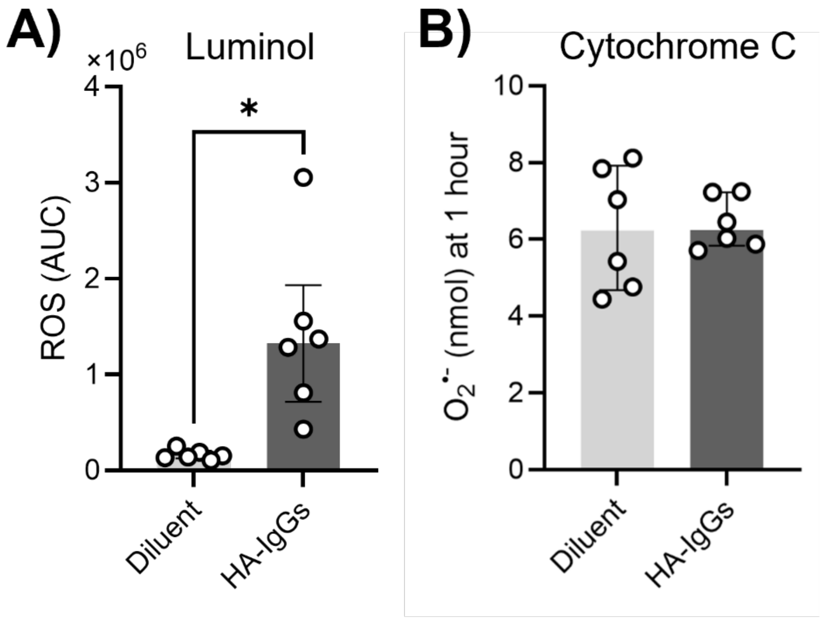


**Supplemental Figure S5. Under serum-free conditions, luminol detects ROS production in response to HA-IgGs. A)** ROS experiment was performed as described in *Method*s, omitting 10% human serum. **B)** Neutrophil-enriched cell suspensions were resuspended at 1 x $10^6$ cells/mL in HBSS supplemented with 125 µM cytochrome C before stimulation with HA-IgGs (1 mg/ml). Optical density was measured at 550 nm (reference 540 nm) and superoxide anion production was calculated as previously described (C. Dahlgren and A. Karlsson. J Immunol Methods 1999 Vol. 232 Issue 1-2 Pages 3-14. PMID: 10618505). Data are shown as individual values from six different donors with median and interquartile range. Statistical analysis was performed using the Wilcoxon signed-rank test ($P < 0.05$). AUC: Area Under the Curve, HA-IgGs: Heat-Aggregated IgGs, ROS: Reactive Oxygen Species.

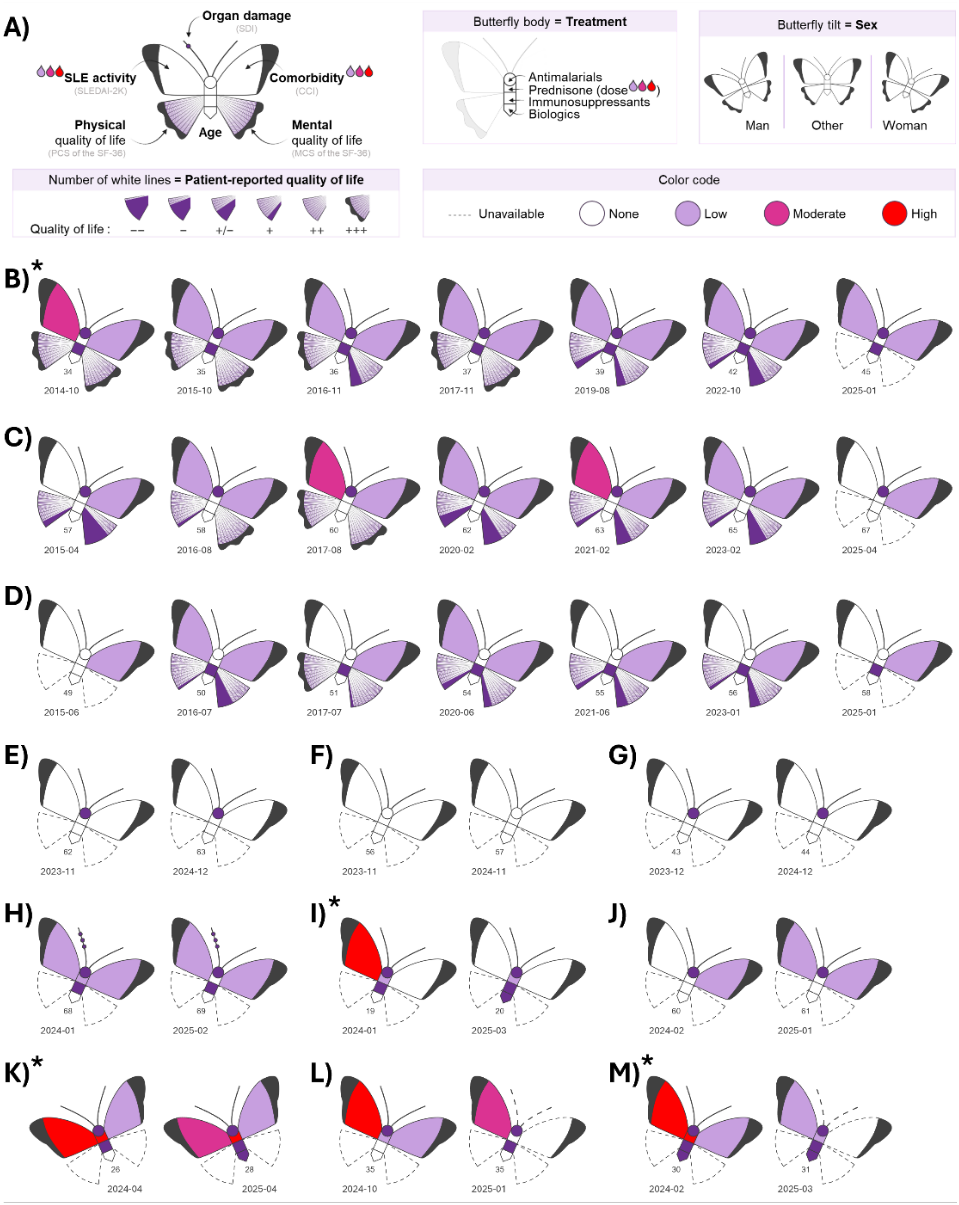


**Supplemental Figure S6. Purple Butterfly series of the 12 SLE patients included in the present study. A)** Guide to interpreting the Purple Butterfly (for a detailed description, please refer to DOI: 10.1093/rap/rkae075). **B-M)** Individual patient trajectories, with each panel representing a single patient. *Patients with lupus nephritis (reported at or before the first illustrated visit). CCI: Charlson Comorbidity Index, MCS: Mental Component Summary, PCS: Physical Component Summary, SDI: SLICC/ACR Damage Index, SLE: Systemic Lupus Erythematosus, SLEDAI-2k: Systemic Lupus Erythematosus Disease Activity Index 2000, SF-36: 36-Item Short-Form Health Survey.
